# Functional ultrasound imaging and neuronal activity: how accurate is the spatiotemporal match?

**DOI:** 10.1101/2024.07.10.602912

**Authors:** Théo Lambert, Hamid Reza Niknejad, Dries Kil, Clément Brunner, Bart Nuttin, Gabriel Montaldo, Alan Urban

## Abstract

Over the last decade, functional ultrasound (fUS) has risen as a critical tool in functional neuroimaging, leveraging hemodynamic changes to infer neural activity indirectly. Recent studies have established a strong correlation between neural spike rates (SR) and functional ultrasound signals. However, understanding their spatial distribution and variability across different brain areas is required to thoroughly interpret fUS signals. In this regard, we conducted simultaneous fUS imaging and Neuropixels recordings during stimulus-evoked activity in awake mice within three regions the visual pathway. Our findings indicate that the temporal dynamics of fUS and SR signals are linearly correlated, though the correlation coefficients vary among visual regions. Conversely, the spatial correlation between the two signals remains consistent across all regions with a spread of approximately 300 micrometers. Finally, we introduce a model that integrates the spatial and temporal components of the fUS signal, allowing for a more accurate interpretation of fUS images.

## Introduction

Functional ultrasound imaging (fUS) is a neuroimaging technique that has been increasingly adopted by the neuroscience community in recent years^1–3^. Its high spatiotemporal resolution and brain-wide capabilities^4–6^, along with its ease of use, have positioned the technology as a valuable tool for probing brain-wide functional networks^6–9^. Nevertheless, fUS does not directly report on neuronal activity but on hemodynamics (mostly cerebral blood volume but also blood velocity^4,5,10^), known to be linked together through the neuro-glio-vascular coupling^11–13^. As many spatio-temporal aspects of this coupling remain elusive at the mesoscale, the interpretation of the fUS signal is not straightforward^14^.

Using separate fUS and electrophysiology recordings, several studies have already demonstrated regional correlation between both types of activity^7,8,15–17^. However, these experiments merely served as control and did not allow for a thorough exploration of the link between fUS and neuronal signals. To gain a deeper understanding of the connection between both signals simultaneous fUS imaging and electrophysiological recording must be performed, a process that is technically challenging. Aydin et al.^18^ paved the way with the sequential use of Ca^2+^ / fUS imaging in anesthetized mice and modelled the temporal response in the olfactory bulb with a gamma function.

These results were complemented by Nunez-Elizalde et al.^19^ who performed simultaneous fUS and high-density electrode array recordings in awake mice, from which they obtained a linear relationship in the <0.3 Hz band and a similar temporal transfer function in the visual cortex and hippocampus. Finally, Claron et al.^20^ reported the temporal correlation of the fUS signal with single unit activity in non-human primates during a behavioral task but not in resting state. These dedicated studies have been incrementally building knowledge regarding the relevance of fUS signals from a neuronal perspective. However, several key aspects were not addressed yet.

First, no study has examined the spatial concordance between neuronal activity and hemodynamic responses; the latter frequently extends beyond the boundaries of the active neuronal network^21–23^. Furthermore, the quantitative relationship between neuronal spike rates and the fUS signal has not been defined. Lastly, whether the correspondence between fUS signals and neuronal activity is uniform throughout the brain or differs among regions remains uncertain^24^.

To address these questions, we used a high-density multi-electrode array, the Neuropixels probe^25^, combined with fUS to simultaneously capture the hemodynamic signal and neuronal activity. Recordings were carried out separately in three distinct regions of the visual pathway: the visual cortex, the superior colliculus and the lateral geniculate nucleus for several stimulus contrasts (0-100%). Each of these regions elicited varying proportionality factors between the fUS amplitude and spike rate. We further examined the spatial and temporal transfer functions between both signals, that were finally summarized in a computational model. From this comparison, our investigation aims to determine how to interpret the fUS signal in terms of neuronal activity.

## Results

### Simultaneous fUS and Neuropixels recordings

Simultaneous and concurrent fUS and Neuropixels recordings were performed in three regions of the visual pathway, namely the superior colliculus (SC), the lateral geniculate nucleus (LGN) and the primary visual area (V1; **Figure 1**.a). The insertion of the Neuropixels probe was guided from preliminary fUS scans to ensure i) optimal colocalization and ii) high reproducibility of fUS and Neuropixels recordings over sessions and animals (**Figure S1**; **Materials and Methods** – Simultaneous fUS and Neuropixels recordings). The probe trajectory was determined in structural B-mode ultrasound images during the retraction (**Figure 1.b**) and the position was estimated *post-hoc* (**Materials and Methods** – Probe trajectory).

**Figure 1.**
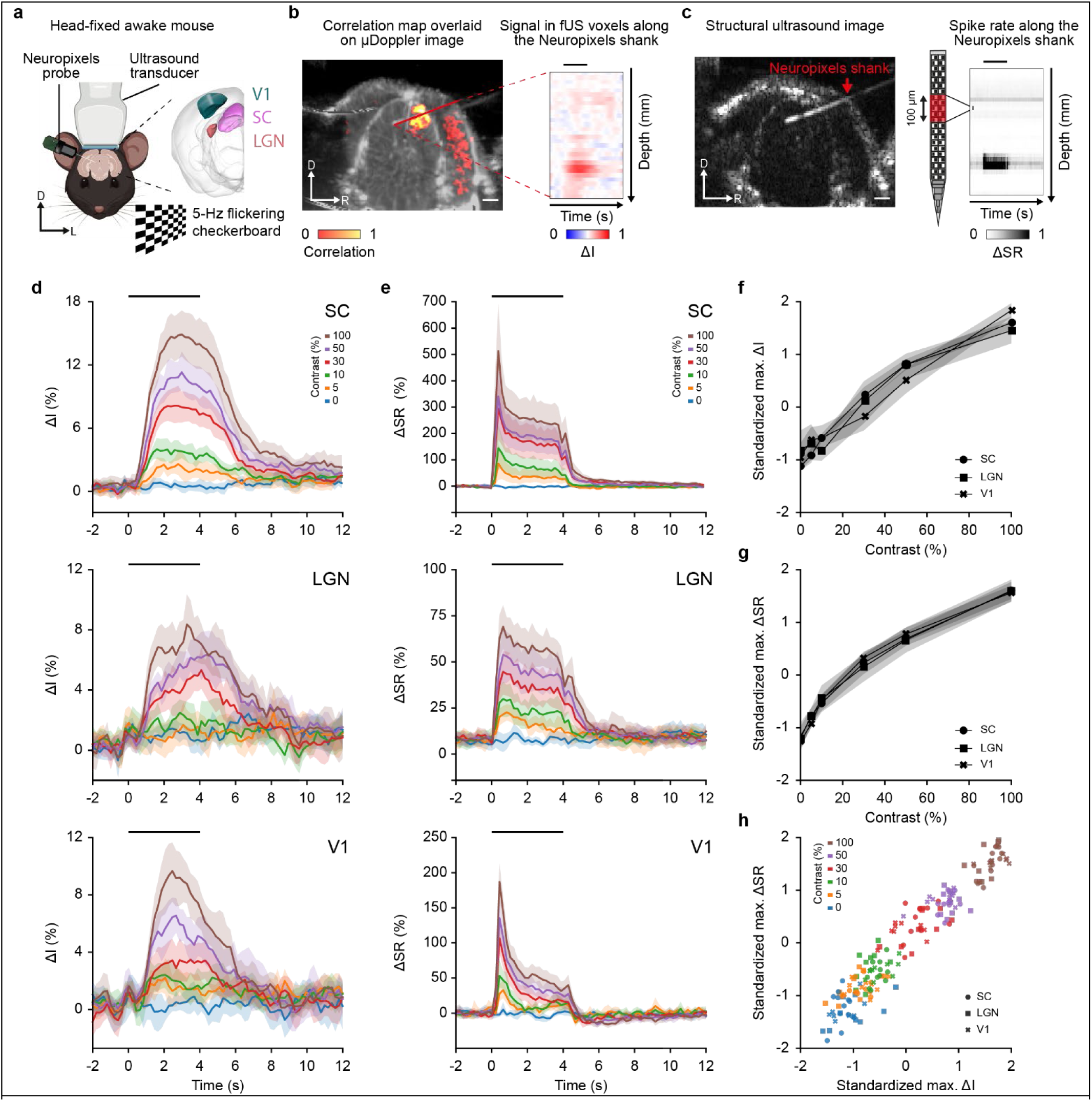
Joint functional ultrasound imaging and Neuropixels recording in the visual pathway. **a.** Schematic representation of the joint fUS-Neuropixels recording setup. Mouse was prepared with a cranial window allowing for combined fUS imaging and Neuropixels recording of either the superior colliculus (SC, pink), the lateral geniculate nucleus (LGN, orange) or the primary visual area (V1, green). Awake mouse was subjected to full screen 5-Hz flickering checkerboard stimuli, with varying contrast, covering the entire visual field of the left eye. Stimuli parameters are provided in the **Materials and Methods**. D, dorsal; L, left. **b.** *Left,* Correlation map extracted from fUS imaging, overlaid with the µDoppler image. *Right,* Example time-traces of change in fUS signal (ΔI in %) extracted from fUS voxels crossing the Neuropixels trajectory, same mouse as (**b**). Example correlation maps for SC, LGN and V1 can be found in **Figure S3**. **c.** *Left,* Structural (B-mode) image of the mouse brain. The echoes backscattered by the Neuropixels probe allows the accurate location of the shank. *Right,* Example of variation in spiking rate (ΔSR) along the Neuropixels shank downsampled to match the fUS resolution (100 µm, 5 Hz). D, dorsal; R, right. For (**b**, **c**), the horizontal black line depicts the stimulation period, and scale bars are 1 mm. D, dorsal; R, right. **d.** Time-course variation in fUS signal (ΔI in %; mean ± 95% CI) in SC, LGN and V1 (top to bottom), extracted by computing the average signal in a band of 3 voxels around the Neuropixels shank. **e.** Time-course variation in spike rate (ΔSR in %; mean ± 95% CI) in SC, LGN and V1 (top to bottom), obtained by averaging the signal in all bins crossing the regions of interest. For (**d**, **e**), the stimulus was presented at time 0 for 4 s and is denoted by the horizontal black line. **f.** Maximum of fUS signal variation (max. ΔI in %) with respect to the contrast conditions, averaged across animals and sessions (mean ± 95% CI). ΔI was standardized per session and region (zero-mean, unit variance) to account for the differences in scale. **g.** Maximum of spike rate variation (max. ΔSR in %) with respect to the contrast conditions, averaged across animals and sessions (mean ± 95% CI). ΔI was standardized per session and region (zero-mean, unit variance) to account for the differences in scale. **h.** Standardized maximum of spike rate variation (max. ΔSR in %) with respect to the standardized maximum of fUS signal variation (max. ΔI in %) across animals and sessions. For (**d, e, h**), contrast conditions: blue 0%, orange 5%, green 10%, red 30%, purple 50%, brown 100%. For (**f-h**), SC: round, LGN: square, V1: cross.

Spiking activity in the three regions of the visual pathway was modulated by a visual stimulus consisting of a 4-s, 5-Hz flickering checkerboard with contrast levels of 0, 5, 10, 30, 50, and 100%, randomized between trials. Multiple recording sessions of the regions of interest were performed in several mice (SC: 8 mice / 12 sessions, LGN: 7 / 9, V1: 5 / 9. See details in **Table S1**). Pupil size measurements confirmed that the mice were awake and alert during the recording sessions (**Figure S2**).

fUS captures a signal proportional to the cerebral blood volume within a full cross-section of the mouse brain with a spatial resolution of ∼100×110×200 µm^3^ ^26^ and a frame rate of 5 Hz (**Materials and Methods** – Functional ultrasound imaging sequence). To allow strict comparison across mice, sessions, regions and contrast conditions, the fUS signal was normalized to the baseline to obtain a relative change in signal intensity, termed ΔI. Functional images were computed by correlating ΔI with the stimulus pattern (Figure 1**.b, left panel**). Correlation maps for each region investigated can be found in **Figure S3**.

On the other hand, the Neuropixels probe recorded electrical activity from the brain surface down to the tip of the probe in a ∼40-µm radius cylinder with a density of 1 electrode per 20 µm^25^. The spike rate (SR) was extracted from the raw data following standard preprocessing pipeline (**Materials and Methods** – Electrophysiological data processing) before being spatially and temporally sampled to match the resolution of the fUS data i.e., 100µm and 5Hz. Finally, the data was normalized to obtain a relative variation in spiking rate (ΔSR) making the comparison across mice, sessions, regions and contrast conditions easier. This procedure allows to depict the ΔSR along the shank and over time (Figure 1**.c**; **Materials and Methods –** Electrophysiological data processing).

To compare the fUS signal with the Neuropixels data, the ΔI was extracted from the voxels crossing the path of the Neuropixels probe (Figure 1**.c., right panel. Materials and Methods** – Probe trajectory). Such an alignment between modalities allows a direct comparison between ΔI and ΔSR in terms of time-course, amplitude, spatial correspondence, and thus transfer function.

### Contrast-dependent fUS signal and spike rate

To compare the temporal shape of the functional responses, we spatially averaged the ΔI and ΔSR along the regions of interest (**Materials and Methods** – Trajectory based region averaging). Hemodynamic responses in SC, LGN, and V1 exhibited a steep increase during the stimulus presentation followed by a long return-to-baseline period (Figure 1**.d** - top to bottom respectively). These responses were detected from the lowest contrast used, i.e., 5% (except for the LGN which was detected as from 10%), and with a constant increase of the response amplitude up to the 100% contrast condition.

The averaged ΔSR shows a peak immediately after stimulus presentation, followed by a slow decrease in spike rate and a fast return-to-baseline period at the end of the stimulation. Like the hemodynamic response, ΔSR follows a positive contrast-response, which is most prominent in V1 and SC, followed by LGN (Figure 1**.e**).

As hemodynamics and spiking responses increase with contrast, the maximum values of ΔI and ΔSR for each contrast were extracted and normalized with respect to their baseline (mean of 0 and unit variance) to account for scaling differences between regions. The maximum of ΔI as a function of contrast follows the same dynamics for SC, LGN and V1 (Figure 1**.f**), with similar results observed for the ΔSR (Figure 1**.g**). A direct comparison of the normalized maxima of ΔI to ΔSR reveals a linear relationship between the two measures for all three recorded regions (Figure 1**.h**).

### Linear relationship between fUS and spike rate

The relationship between ΔI and ΔSR amplitudes is a first element that allows a direct interpretation of the fUS signal in terms of spike rate. To investigate further, we compared the maximum of ΔI and ΔSR for each probe insertion and observed a linear fit between the two measurements for each contrast condition (dots), session (color-coded), and region (SC, LGN, and V1; left-to-right in Figure 2**.a**). Non-normalized data are presented in **Figure S4**. This was further confirmed by computing the coefficients of determination per session which reached 0.94 ± 0.04 for SC, 0.89 ± 0.06 for LGN and 0.92 ± 0.03 for V1 (mean ± sd; Figure 2**.b**). Note that these results do also stand for the area under the curve value associated to ΔI since the two are highly correlated (R^2^>0.99; **Figure S5**).

**Figure 2.**
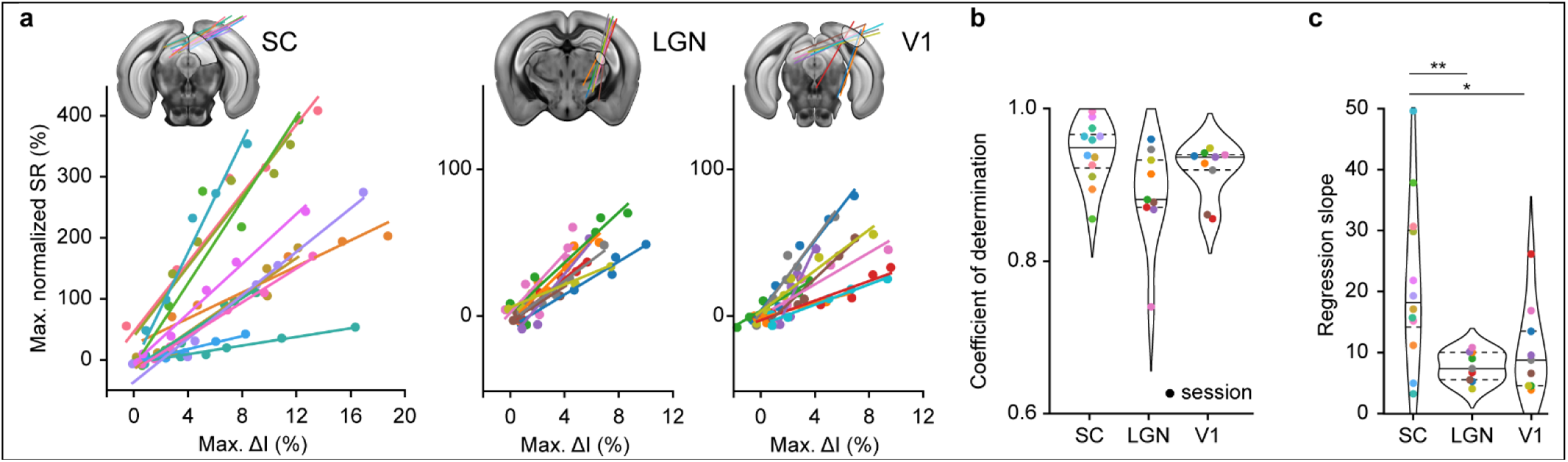
Linear relationship between hemodynamic amplitude (fUS) and spike rate (Neuropixels). **a.** Normalized maximum of spike rate variation (ΔSR in %) with respect to the normalized maximum of fUS amplitude (max. ΔI in %). Color corresponds to a recording session and dot represents a contrast condition. ΔSR is defined as the SR divided by the average SR during baseline in the bins located in the region of interest. Neuropixels trajectories are overlaid with reference atlas (CCF v3) for the superior colliculus (SC), the lateral geniculate nucleus (LGN) and the primary Visual area (V1), from left to right. Non-normalized data are shown in **Figure S4**. Example of Neuropixels probe trajectories can be found in **Figure S6**. **b.** Distribution of the coefficients of determination in SC, LGN, and V1. Colored dot corresponds to a recording session. Horizontal lines indicate first/third quartiles (dotted) and median (plain line). **c.** Distribution of the regression slopes in SC, LGN, and V1. Colored dot corresponds to a recording session. Horizontal lines indicate first/third quartiles (dotted) and median (plain line). P-values: *p < 0.05, **p < 0.01, t-test of independent samples. The color code is shared across a-to-c panels.

Given the high linearity between the amplitudes of ΔI and ΔSR we examined the slopes, that provide a calibration between the two measures. Their values are 21.4 ± 13.0 for SC, 7.7 ± 2.3 for LGN, and 10.5 ± 6.9 for V1 (mean ± sd; Figure 2**.c**). Such slopes correspond to the relative change in spike rate associated to 1% increase in the fUS signal maximum amplitude. Interestingly, the distribution of slopes in the SC significantly differs from those in the LGN (t-test, p<0.01) and V1 (t-test, p<0.05), suggesting that the relationship between ΔI and ΔSR is not universal across the regions of the visual pathway, and thus the brain.

### Differences in temporal transfer functions across brain regions

To model the temporal link between SR and fUS responses, we assumed the existence of a causal and linear relationship and followed the procedure described in Nunez-Elizalde et al.^19^. The transfer functions (TF) were calculated using a linear regression and convolved with the SR to estimate the associated fUS responses (Figure 3**.a**; **Materials and Methods** – Transfer function estimation).

**Figure 3.**
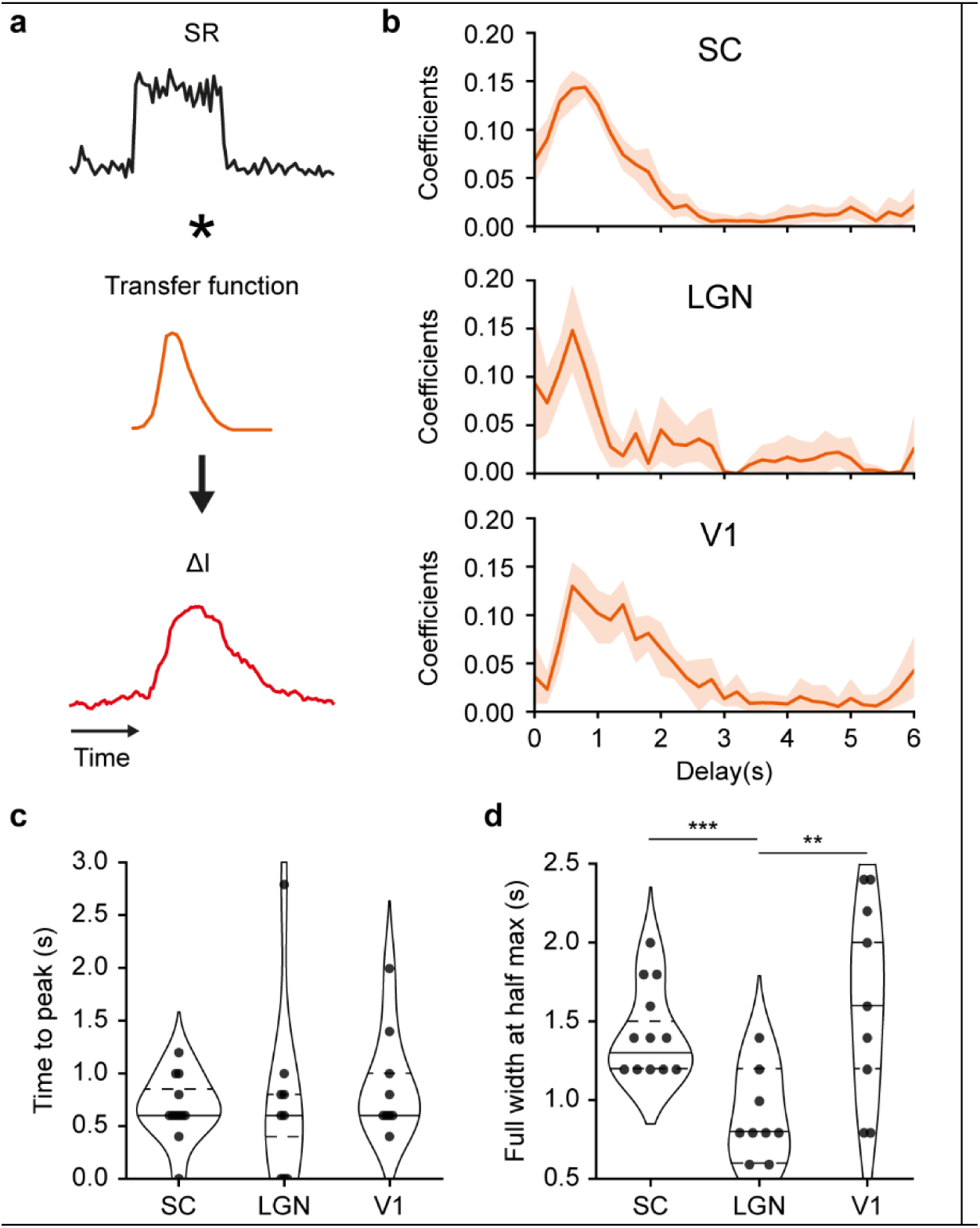
Temporal transfer of spike rate to fUS signals. **a.** Model employed to convolve the spike rate (SR; black curve) as a transfer function (TF; orange curve) to obtain a temporal trace resembling the fUS signal (ΔI; red curve). The TF is estimated through linear ridge regression. **b.** Average temporal transfer functions (TF; mean ± 95% CI) computed for the superior colliculus (SC), the lateral geniculate nucleus (LGN) and the primary visual area (V1; top to bottom). Coefficients are referring to the parameters of the linear regression. TF were estimated for each session individually. **c.** Distribution of the TF’s time-to-peak values (s) across sessions (dots) and for SC, LGN and V1 regions. **d.** Distribution of the temporal TF’s full width at half maximum values (ΔFWHM in s) across sessions (dots) and for SC, LGN and V1 regions. Horizontal lines indicate first/third quartiles (dotted) and median (plain line). P-values: ** p<0.01; ***p<0.001, t-test of independent samples.

The estimated transfer functions for the 3 regions follow a gamma-like shape (Figure 3**.b**) as reported in the literature^18,19^. These TFs show an average time to peak of 0.8 ± 0.5 s (mean ± sd), which is similar across the 3 regions assessed in this study (Figure 3**.c**). However, the estimated TFs differ in their full-width at half maximum, which is shorter for the LGN (0.8 ± 0.3 s; mean ± sd) when compared to SC (1.5 ± 0.3 s, ***p<0.001, t-test) and V1 (1.5 ± 0.6 s, **p<0.01, t-test; Figure 3**.d**). These temporal differences suggest that the transfer function between ΔI and ΔSR is not universal across the brain.

### Accurate spatial match between fUS signal and spike rate

We examined the spatial relationship between the fUS signal and spiking activity by comparing their maxima at the highest contrast condition (Figure 4**.a**). The result is presented for the SC, LGN, and V1 in Figure 4**.b-d** respectively, per session, and averaged after peak alignment to assess the spread of the signals. Despite the diversity in peak location (due to different Neuropixels probe locations) and width, both maximum of ΔI and ΔSR showed remarkable alignment, as confirmed by the strong correlation between maximum ΔI and ΔSR curves in all sessions, and regardless of the region of interest (SC: r^2^ = 0.72 ± 0.21; LGN: r^2^= 0.69 ± 0.10; V1: r^2^= 0.73 ± 0.15, mean ± sd; Figure 4**.e**).

**Figure 4.**
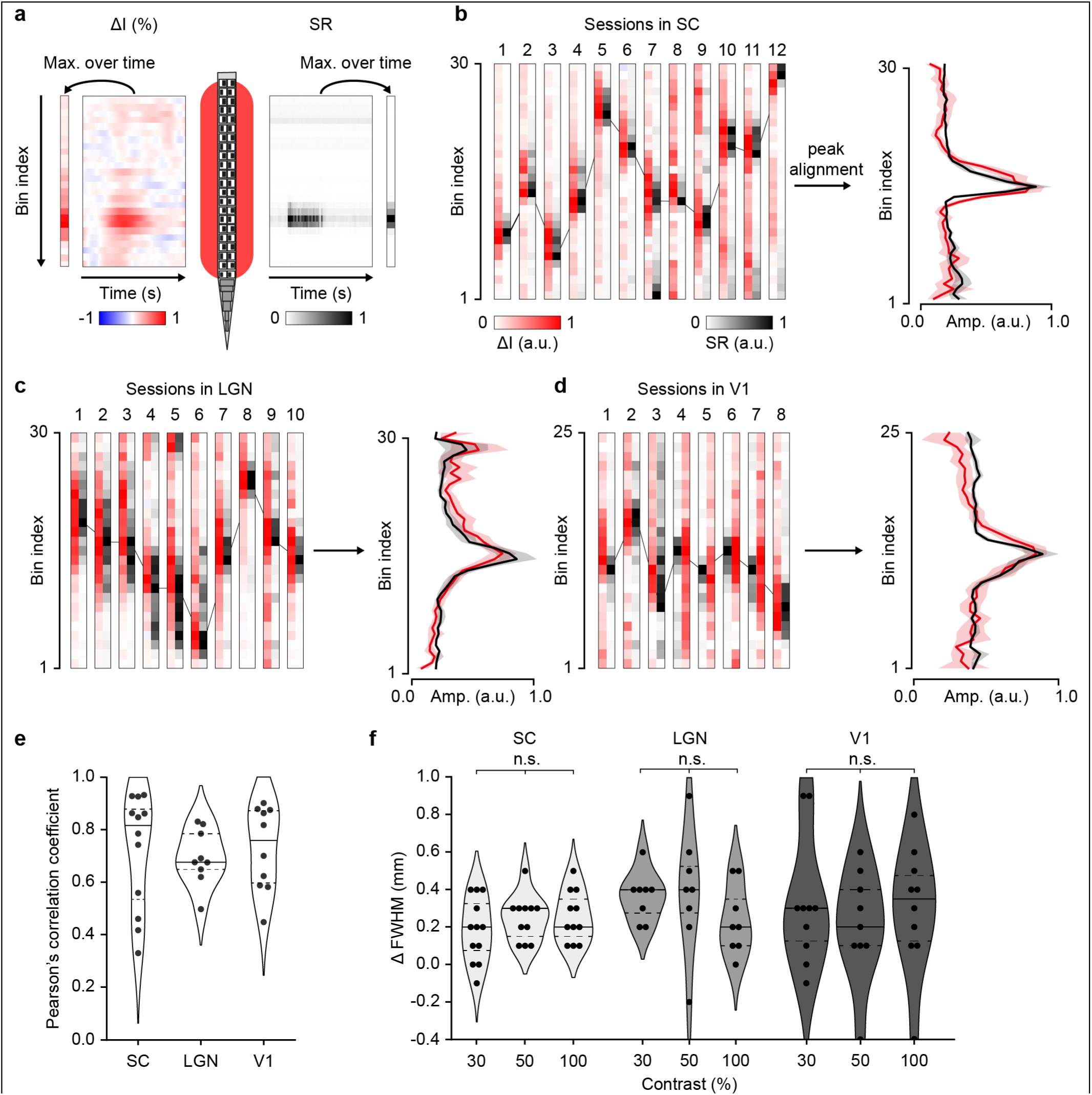
Spatial transfer of spike rate to fUS signals. **a.** Schematic representation of the spatial comparison process between spike rate (SR; Neuropixels schematic) and fUS voxels (red area). fUS signal (ΔI as arbitrary unit, a.u.; left) and spike rate (ΔSR as arbitrary unit, a.u.; right) are temporally reduced by computing their maximum amplitude during the stimulus presentation. **b-d.** Normalized spatial amplitudes of the fUS and spike rate (arbitrary units; a.u.) at 100% contrast in the SC (b), LGN (c), V1 (d), presented across individual sessions (left) and averaged after peak alignment (mean ± sd; right). **e.** Distribution of Pearson’s correlation coefficients between spatial fUS and spike rate amplitudes, across recording sessions and regions. **f.** Distribution of the spatial TF’s full width at half maximum differences (ΔFWHM in mm) between fUS and spike rate amplitudes, across recording sessions, contrasts, and regions. Horizontal lines indicate first/third quartiles (dotted) and median (plain line). A positive difference indicates a wider spatial extent of the fUS signal as compared to the spike rate. Each point represents a session.

Finally, to measure the spatial spread of the fUS signal, we computed and compared the spatial full width at half maximum in the aligned ΔI and ΔSR (Figure 4**.f**). Such analysis was performed at 30, 50 and 100% contrast since sufficient signal-to-noise is required to accurately delineate the active area. We observed a larger response of ∼0.3 mm (∼3 fUS voxels) for the hemodynamic signal (ΔI), which was comparable across regions (0.28 ± 0.16, 0.33 ± 0.32, 0.30 ± 0.31 mm at 100% contrast in SC, LGN and V1, respectively). Importantly, the values did not significantly vary with the contrast, suggesting that higher neuronal responses do not induce larger spread (Kruskal-Wallis test, SC: p=0.56, LGN: p=0.26, V1: p=0.78).

### Spatiotemporal model

Based on the results of our study and existing literature, we propose a model of the relationship between ΔSR and ΔI. According to previous research, the spatiotemporal TF can be considered as two convolutions, with a spatial (𝑔) and a temporal (ℎ) TF. We demonstrated above that the spatial spread can be approximated to 0.3 mm in all the regions included in our study. Therefore, 𝑔 is independent of time and can be modeled as a Gaussian filter 𝐺(𝑥; µ, 𝜎) with zero mean and standard deviation of 0.15 mm:

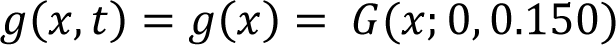

In line with prior work^18,19^, we show that utilizing a gamma-like linear filter for the convolution provides a satisfactory approximation of ℎ. Nevertheless, some parameters of such filter vary across brain regions. Thus, with 𝛤 denoting the gamma function and 𝜃_𝑟_ the parameters associated with a region:

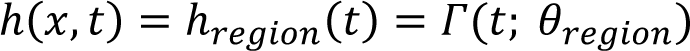

Merging the equations produces the final model of the relationship, where spatial and temporal convolutions are denoted by ⊗_𝑥_ and ⊗_𝑡_, respectively:

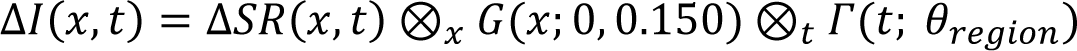

## Discussion

In this work we performed simultaneous fUS imaging and Neuropixels recording in three regions of the visual pathway in awake mice. We demonstrated that regardless of the regions investigated, the fUS signal (i.e., cerebral blood volume) is linearly related to the local neuronal spiking activity with a limited spatial spread. Moreover, we found consistent regional differences in the proportionality coefficients and in the duration of the transfer functions. Finally, we proposed a spatiotemporal theoretical model that incorporates these findings. Based on these results, we discuss the interpretability of the fUS signal in terms of spiking activity.

The linearity between fUS and spike rate amplitudes indicates that a relative change in spike rate can be accurately related to a similar change in fUS signals. However, a calibration between ΔI and ΔSR remains difficult as the proportionality coefficients showed a large variability within the region of interest and significant differences between regions (SC: 21.4 ± 13.0, LGN: 7.7 ± 2.3, V1: 10.5 ± 6.9; mean ± sd).

The variability within the same region might be attributed to several factors: insertion site, number of recorded units, eye position of the animal during the stimulus presentation, arousal, etc. On the other hand, the difference between the regions can be explained by intrinsic disparities in the brain such as in cell diversity^27^, stimulus configuration^28^, vascular architecture^29^ and neurovascular coupling mechanisms^24^. Note that the normalization of the spike rate (ΔSR) is independent of the number of units recorded but relies on its basal activity. In the case of the SC, the latter is very low thus resulting in large ΔSR values.

The linear relationship with spiking activity was validated for fUS amplitudes ranging from 0 to +20%, therefore some cases such as negative or higher amplitude signals are not addressed in this study. Recently, Macé et al.^7^ reported a negative fUS signal correlated with a decrease in basal spiking activity. It remains uncertain whether the same linearity coefficient and filter can be applied for positive and negative fUS signals. On the other extremum, high fUS increases of ∼50% have been found during spreading depolarizations^4,30,31^. In such pathological conditions, the neurovascular coupling is largely disrupted, requiring specific interpretation of the link between spiking activity and fUS signal^11^.

Regarding the temporal filter, the time-to-peak values were independent of the region but some significant differences in the length of the transfer function (∼1 s) were observed. Without calibration, the comparison of fUS signals’ durations between regions must be carefully conducted, as one cannot determine whether the differences are due to different spiking activity or to a different transfer function.

For the spatial aspect, we observed a limited spread (300 µm) of the fUS signal as compared to the spiking counterpart, regardless of the contrast value. This finding suggests that fUS images displaying small, activated regions or small changes in the size of the active area can be interpreted with high reliability.

Our experiments were performed in the visual pathway allowing for the simple modulation of spiking activity in three distinct regions located either in the thalamus (LGN), the midbrain (SC) or the cortex (V1). Given that our results were comparable to those in the olfactory bulb^18^, it is reasonable to expect that they will generalize well to other sensory systems. However, a comprehensive study of the whole brain may be very challenging as arbitrarily stimulating a given region is not a straightforward task. Different approaches, such as using resting state activity or optogenetic stimulation, may offer a solution to achieving a complete mapping of the relationship between fUS and spiking activity^8,19^.

Given the current evidence, we argue that fUS images can be interpreted as a robust estimator of local spiking activity because both signals are linearly correlated with minimal spatial spread. Local fUS signal variations in amplitude, spatial extent, or temporal shape then have a proportional counterpart in the spike rates. Therefore, fUS is a highly precise map-making tool that can inform neuroscientists at the whole-brain level on where to look for mechanistic explanations with complementary modalities. Furthermore, combined with existing solutions for neuromodulation^32^, our findings establish a path to achieve dependable full ultrasound control of neuroprosthetics^33,34^.

## Materials and methods

### Animals

All experimental procedures were approved by the Ethical Committee for Animal Experimentation of the KU Leuven. Twelve male adult C57BL/6 mice (2-4 months old; Janvier Labs) were used for the experiments, details are provided in **Table S1**. Prior to surgery, mice were kept in group cages in a 12-hr dark-light cycle environment at a constant temperature of 21°C. Prior to the experimental procedure, mice were handled daily and habituated to non-aversive hand-cupping. Once accustomed to the experimenter (e.g., exhibiting grooming behaviors), mice are progressively trained to walk through the body tube of the experimental setup. After surgery, mice were individually housed in an enriched cage (nesting material, shelter, wood block) with *ad libitum* access to food and water. All experiments were conducted during the light phase of the mouse’s activity cycles.

### Surgical procedure

The surgical procedure closely follows a published protocol^26^. At the start of the procedure, anesthesia was induced with an intraperitoneal injection of mix of Ketamine (100 mg/kg, Nimatek, Dechra) and Medetomidine (1 mg/kg, Dormitor, Orion Pharma). The paw of the animal was pinched regularly to check for the absence of a pedal reflex. Once deeply anesthetized the mouse was fixed in a stereotactic frame equipped with a homeothermic blanket to maintain a stable body temperature around 36.5°C. Eye ointment was applied to prevent dehydration of the conjunctivae (Duratears, Novartis). Subsequently, the head was shaved and cleaned (Iso-betadine, Meda) and the scalp was incised and removed to expose the dorsal part of the skull. Lateral and posterior muscles were disconnected and retracted. A stainless-steel headplate was attached to the skull using dental cement (Superbond C&B, Sun Medical). Then a cranial window extending from Bregma 0 to -6 mm and ±4.5 mm apart from the sagittal suture was performed using a high-speed rotary hand drill (Foredom). Once the brain was exposed and cleaned with sterile saline, the cranial window was protected with a layer of silicone elastomer (Body Double-Fast Set, Smooth-on). At the end of the procedure, the mouse was allowed to recover for 24 hrs in its homecage placed on a heating pad, followed by 7 days of recovery during which the mice was treated with painkillers (0.2 mg/kg i.p., Buprenorphine, Vetergesic, Ecuphar) antibiotics (15 mg/kg i.p., Cefazolin, Sandoz; 1% Emdotrim in drinking water, Ecuphar), and anti-inflammatory drugs (0.1 mg/kg Dexamethasone, Rapidexon, Dechra) during the first three days. Dexamethasone and Emdotrim were continued for a total of 5 days post-surgery.

### Habituation to head-fixation

Once they have recovered from the surgical procedure, implanted mice were daily habituated to head-fixation after walking through the body tube^26^. Fixation periods were progressively increased, from 5 mins the first day up to 3 hrs after two weeks. A 5% sucrose solution (Sigma-Aldrich, USA) was used to reward the mouse during the habituation sessions.

### Visual stimulus

Visual stimuli were presented on a 32-inch LCD monitor (Samsung S32E590C, 1920×1080 pixel resolution, 60-Hz refresh rate, average luminance of 2.6 cd/m^2^) positioned at an angle of 45° to the mouse head, 20 cm from the left eye, so that the screen was covering 100° of azimuth and 70° of elevation of the left visual field. Each trial consisted in the presentation of a 10-s baseline period with a grey background (50% luminance), followed by a 4-s full screen flickering checkerboard (5-Hz frequency, 0.05 cycles/degree spatial frequency), and ending with a 20-s post-stimulation period with a grey background (50% luminance). The contrast of the checkerboard was set at 0, 5, 10, 30, 50, or 100%. Each session consisted of 300 trials - 50 per contrast condition - presented randomly.

### Functional ultrasound imaging sequence

The imaging procedure used during the experiments was adapted from the sequence for fast, whole-brain fUS imaging previously described^7,26^. The ultrasound transducer containing a linear array of 128 piezoelectric elements was used to emit plane waves (15 MHz, L22-14v, Vermon) in five different angles (−6°, -3°, 0°, 3°, 6°; 3 averages per angle, 500-Hz rate). The backscattered echoes were received and adjusted with a time-gain compensation to account for the attenuation of the signals with increasing depth (exponential amplification of 1 dB/mm). The resulting images were combined to create a high-quality compound image every 200 ms (5 Hz). When the ultrasound waves passed through blood vessels, which contain moving red blood cells, the successive backscattered echoes were phase shifted (Doppler effect), proportional to the speed with which the red blood cells were moving. These shifts were measured and extracted in real-time, using singular-value-decomposition (SVD)-based spatiotemporal filtering, and high-pass temporal filtering (cut-off frequency: 20 Hz). By computing the mean intensity of this filtered signal, we get a ‘Power Doppler’ value, which is proportional to the number of red blood cells in a voxel, and hence to the local cerebral blood volume.

The intensity value of a voxel at a given time was calculated as 𝐼(𝑥, 𝑦) = 𝐴(𝑥, 𝑦, 𝑡)^2^ , where 𝐼 is the Power Doppler Intensity; 𝑥,𝑦 are the coordinates of a given voxel in a given plane; 𝐴 is the amplitude of the compound images after filtering and 𝑡 is time. The resulting functional ultrasound image was 143 x 128 voxels in size (H x W), and each voxel measured 80 x 100 x 300 μm^3^. The script employed for fUS acquisition is described and available in Brunner et al., 2021^26^.

### Procedure for simultaneous fUS-Neuropixels recordings

The awake mouse was head-fixed on the recording platform and the silicone cap was carefully removed (after aseptic cleaning) to expose the brain. In the absence of bone landmarks (i.e., Bregma, Lambda), we used the caudal confluence of sinuses as reference to locate the recording target (**Figure S1**).

The location of the superior colliculus (SC), lateral geniculate nucleus (LGN) and primary visual cortex (V1) were identified for each mouse by performing functional scans (coronal orientation, 250-µm step size) using 4-s 5-Hz flickering checkerboard at 100% contrast (10 trials per position). For each joint fUS-Neuropixels recording session, such functional scans were used to determine the sites of recording by positioning the ultrasound transducer over the middle of the region of interest. Hereafter the ultrasound transducer was moved away, and the cranial window was cleaned from gel and agarose to perform a small duratomy.

The Neuropixels probe (v1.0, Imec)^25^, priorly stained with DiI (ThermoFisher), was secured to a holder fixed to a high-precision micromanipulator. The recording site was reached from the caudal confluence of sinuses (zero reference) and the probe inserted at 5 µm/s. All probe trajectories and insertion depths are displayed in **Table S2**.

Once the Neuropixels probe was inserted, the cranial window was filled up with 2% low-gelling agarose (Sigma-Aldrich) and acoustic gel (Aquasonic Clear, Parker Laboratories) that ensures the acoustic coupling between the brain and the ultrasound transducer. The acoustic gel was dyed black to prevent artifact in the data caused by ambient/stimulation light. The ultrasound transducer was oriented in the coronal plane and tilted 15° to the vertical axis allowing for simultaneous and concurrent Neuropixels recordings. The transducer was lowered down and secured couple of mm from the brain surface. The presence of the Neuropixels probe was confirmed on the ultrasound image. A schematic description of the procedure is shown in **Figure S5**.

### Experimental design

fUS images were acquired at 5 Hz for 34 s (170 µDoppler images). Neuropixels data were recorded at a raw sampling frequency of 30 kHz. The fUS and Neuropixels recordings were synchronized using TTL pulses which tagged the trial onset and the start of the stimulus. Each recording session lasted < 3 hrs and consisted of 300 trials - 50 per contrast condition - presented randomly. At the end of the session, the Neuropixels probe was retracted at ∼0.1 mm/s, while structural ultrasound images (B-mode) were acquired for tracking the position. Finally, the cranial window was cleaned from gel and agarose, and the brain covered with silicone. The mouse was allowed to recover for a minimum of 48 hrs before the next recording session.

### Registration and segmentation

µDoppler images were manually aligned to the Allen mouse Brain Common Coordinates Framework v3.0 (CCF)^35^ using a dedicated interface. The µDoppler image was aligned to the reference atlas using morphological features and landmarks (curvature of the brain, large vessels, ventricles, hippocampus). The coronal cross-section was then interpolated to the atlas resolution (50 µm^3^) by nearest neighbor interpolation. Translations and scaling in the 𝑥, 𝑦 axes and in-plane rotations were then estimated and stored in a transformation matrix. This matrix was finally used to rigidly transform the data at each time point, yielding Allen CCF labels for each fUS voxel. Tools used and procedures followed have been previously described^26^.

### Probe trajectories and spatial alignment

At the end of each experimental session, we captured a 5 Hz B-mode ultrasound video of the Neuropixels probe extraction. From this video, we made an initial estimate of the probe trajectory using the approximate insertion angle and tip location. We considered the initial extraction as an estimate for the following reasons: i) the echoes backscattered by the shank were approximately three voxels wide, which made them wider than the shank itself, ii) spatial binning can cause jitter, iii) the probe tip could be situated off the imaging plane.

To correct for these potential errors, we performed a Monte Carlo sampling procedure to determine the optimal alignment between spike rates and fUS voxels that intersect the probe. We randomly sampled 10,000 trajectories defined by:

● a tip (𝑥, 𝑦), with 𝑥 = 𝑥_0_ + 𝛿, 𝑦 = 𝑦_0_ + 𝜃 where (𝑥_0_, 𝑦_0_) denotes the initial tip position estimate, and (𝛿, 𝜃) ∈ ⟦−5,5⟧ two randomly sampled integers corresponding to a shift in terms of voxels,
● a slope 𝛼 = 𝛼_0_ + 𝜀, where 𝛼_0_ is the initial slope estimate and 𝜀 ∈ [0.2, 0.2] is a randomly sampled real value.

Using each trajectory, we obtained 𝑁 sets of integer values that correspond to the coordinates of voxels in the fUS image (SC: 𝑁 = 30, LGN & V1: 𝑁 = 25). It is noteworthy that for recordings involving both V1 and SC, the estimation in V1 was made using an artificial tip of the probe positioned after the end of SC. We defined our final estimate, 𝑇, by maximizing the Pearson’s correlation coefficient between the fUS voxels extracted from 𝑇 and the first 𝑁 Neuropixels recordings’ bins starting from the tip.

### Trajectory-based region averaging

For the fUS signals, all voxels overlapping with both the Neuropixels trajectory (extended with 3 voxels on each side) and the region of interest (e.g., the SC) were averaged to produce a single temporal trace. This yielded one trace per contrast (total of 6 per session and region). Similarly, all bins in the Neuropixels trajectory belonging to the region of interest were averaged, producing a total of 6 temporal traces.

### Electrophysiological data processing

The electrophysiological recordings were spike-sorted using kilosort2^36^ and manually curated with Phy (github.com/cortex-lab/phy). Data curation aimed to identify clusters corresponding to single- and multi-unit activities and remove noisy clusters based on inter-spike interval, autocorrelation, and waveform shape. The data was binned for every 5 contact points to match the fUS voxel size. Likewise, the trace was binned at 200 ms intervals to match the temporal resolution of fUS signals.

### Transfer function estimation

The transfer functions were estimated using the same procedure as that described by Nunez-Elizalde et al.^19^. The process described in **Trajectory-based region averaging** was applied. A matrix 𝑋 was created by stacking Neuropixels data of all contrast conditions between 𝑡 − 20 and 𝑡 for each time 𝑡 (in frames, 1 frame = 0.2s). Similarly, a column matrix 𝑦 was constructed by gathering fUS data at time 𝑡 for all contrasts. Using 𝑋 as input and 𝑦 as output, a linear ridge regression model was applied to these matrices. The transfer function was defined by the coefficients obtained from the model.

### Parameters calculations

We use the following parameters to quantify spatial or temporal properties of the signals in this manuscript:

- Time to peak: the time point associated with the first inflexion point of the signal after stimulus onset.
- Full width at half maximum (FWHM): length of the longest sequence of amplitude values above half the peak amplitude. Outliers were removed by applying the threshold 𝑚 + 2 × 𝑠.

### Pupil diameter tracking

A near infrared camera (50-Hz imaging frequency; Mako G030, Allied Vision), along with an infrared LED array, was used to record the mouse’s face during the experimental sessions. Pupil diameter was estimated with DeepLabCut^37^ using i) multiple pairs of points around the pupil, and ii) fixed points on left and right canthus. The pupil diameter was computed as the median of the distance between all pairs of points around the pupil, normalized by the distance between canthi.

### Brain tissue processing and Neuropixels probe trajectory confirmation

Mice were euthanized 4 weeks after the first recording. A lethal injection of pentobarbital (100 mg/kg i.p. Dolethal, Vetoquinol) was administered. Mice were transcardially perfused with PBS and 4% PFA (Sigma-Aldrich) using a peristaltic pump. Brains were collected and post-fixed overnight in PFA. 100-µm coronal brain slices were prepared using a vibratome (VT1000S, Leica Microsystems) and analyzed with DAPI staining (diamidino-2-phenylindole, ThermoFisher). Coronal brain slices were mounted with DPX mounting medium (Sigma-Aldrich) and scanned with a confocal microscope (LSM900, Leica) to reveal DiI trace of the Neuropixels trajectories.

### Statistics

The information regarding the statistical tests that were used are presented along with the corresponding results and the captions of the associated figures. The tests were performed using the ‘scipy.stats’ package (version 1.7.3, Python 3.8).

## Acknowledgments

This work was supported by grants from Europe: ERANET-NEURON (G0G3721N-UNSCRAMBLY) and MSCA training network (SOPRANI-GAP-101119916), from Belgium: FWO senior research grants (G079623N-TRPM3 and G0C9923N-USNI4TBI) and FWO senior postdoctoral fellowship (12D7523N-BRAINPAIN for CB) and from USA: NIH (R01NS129836-01A1). This research also received funding from IMEC, the Flemish regional government (AI Research Program) and Internal Fund KU Leuven C14/18/099. We thank the NERF animal caretakers including I. Eyckmans, F. Ooms, and S. Luijten, for their help managing the animals.

## Author contributions

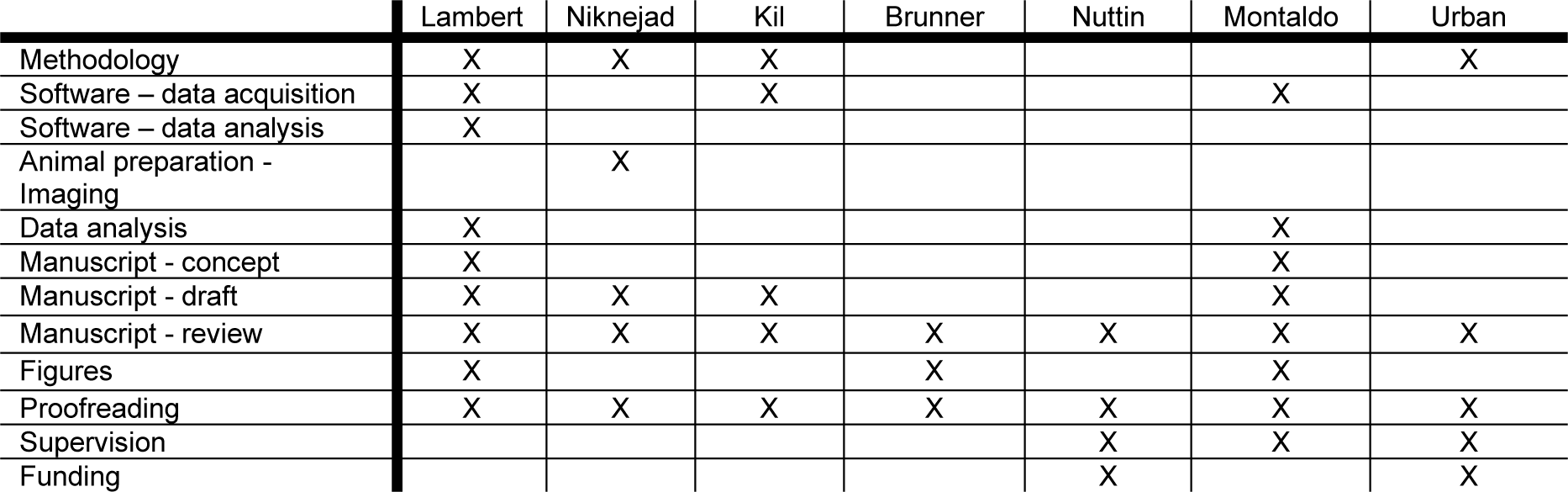

## Supplementary Materials

**Figure S1.**
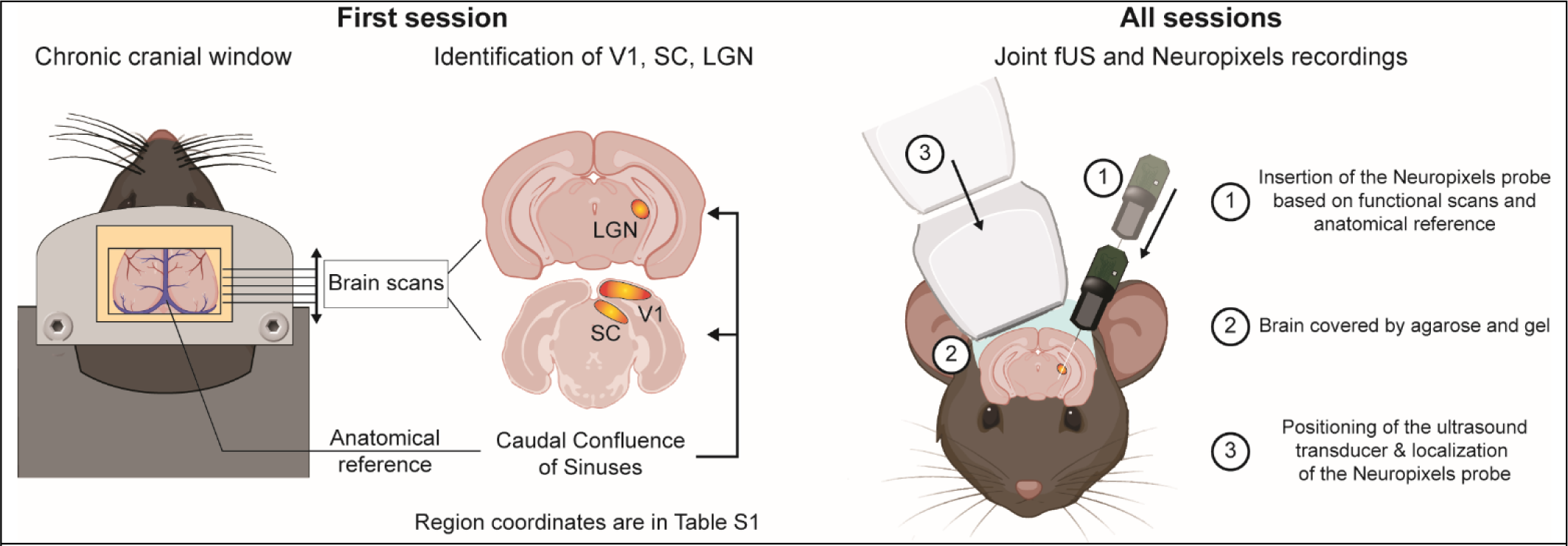
Schematic representation of the data acquisition procedure.

**Figure S2.**
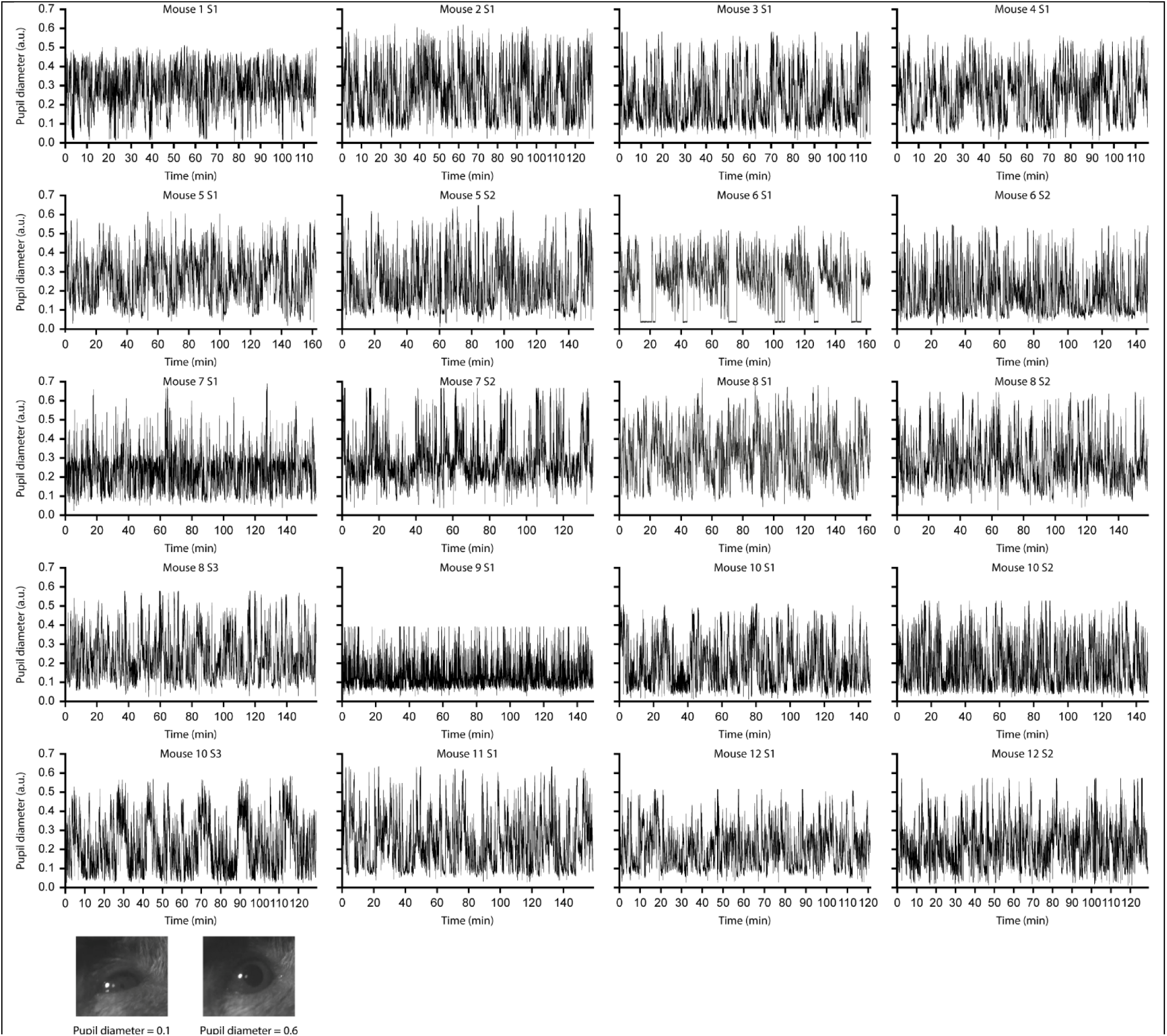
Pupil diameter variation across experimental sessions. *Top,* Normalized variation of pupil diameter tracked across sessions confirming for alertness of mice along recordings. *Bottom,* Example video frames used to compute the mouse’s pupil diameter with small (left) and large pupil size (right).

**Figure S3.**
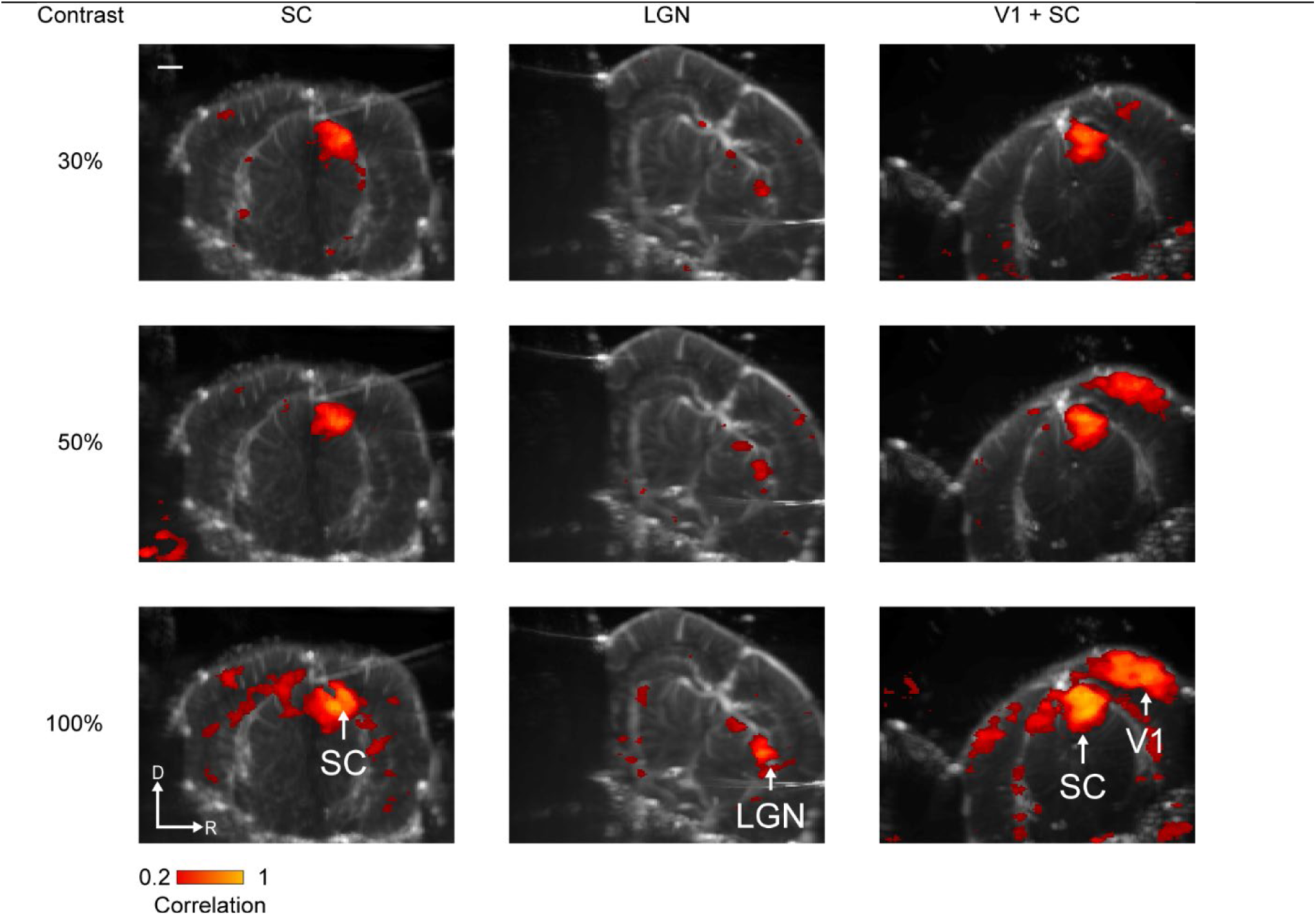
Correlation maps across regions and sessions. Example of correlation maps generated by computing the correlation between fUS voxels and the stimulation pattern across contrast conditions (50 trials each) and regions of interest (one session per region). Scale bar: 1 mm.

**Figure S4.**
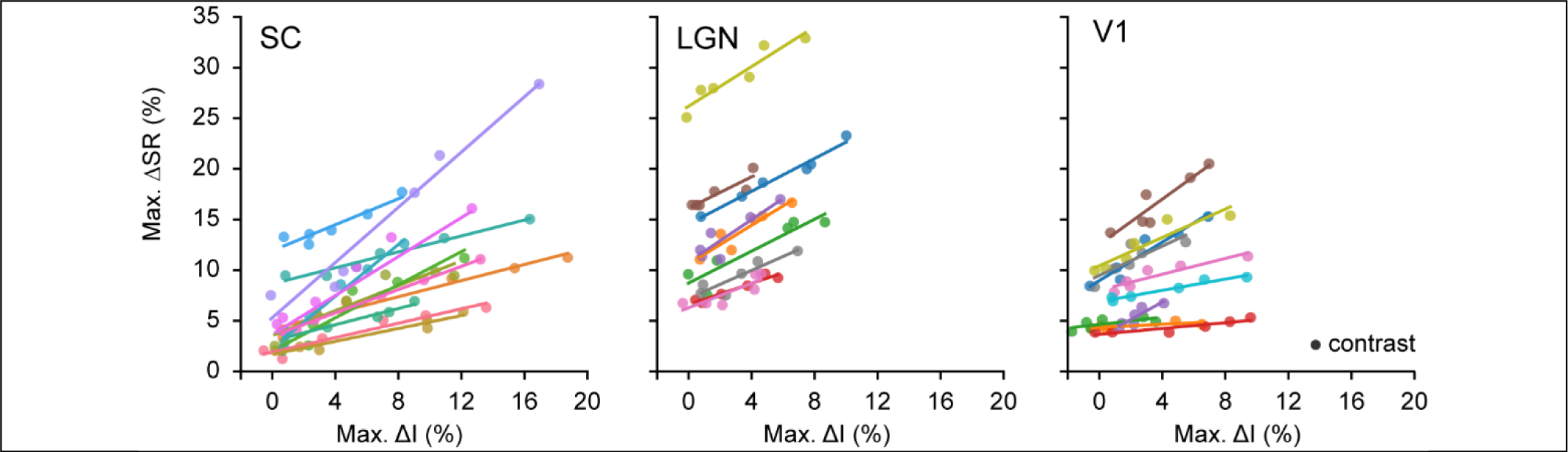
fUS and spike rate activities in responses to contrast conditions. Non-normalized maximum of spike rate variation (max. ΔSR in %) with respect to the maximum of fUS signal variation (max. ΔI in %). Color corresponds to a recording session and dot represents a contrast condition.

**Figure S5.**
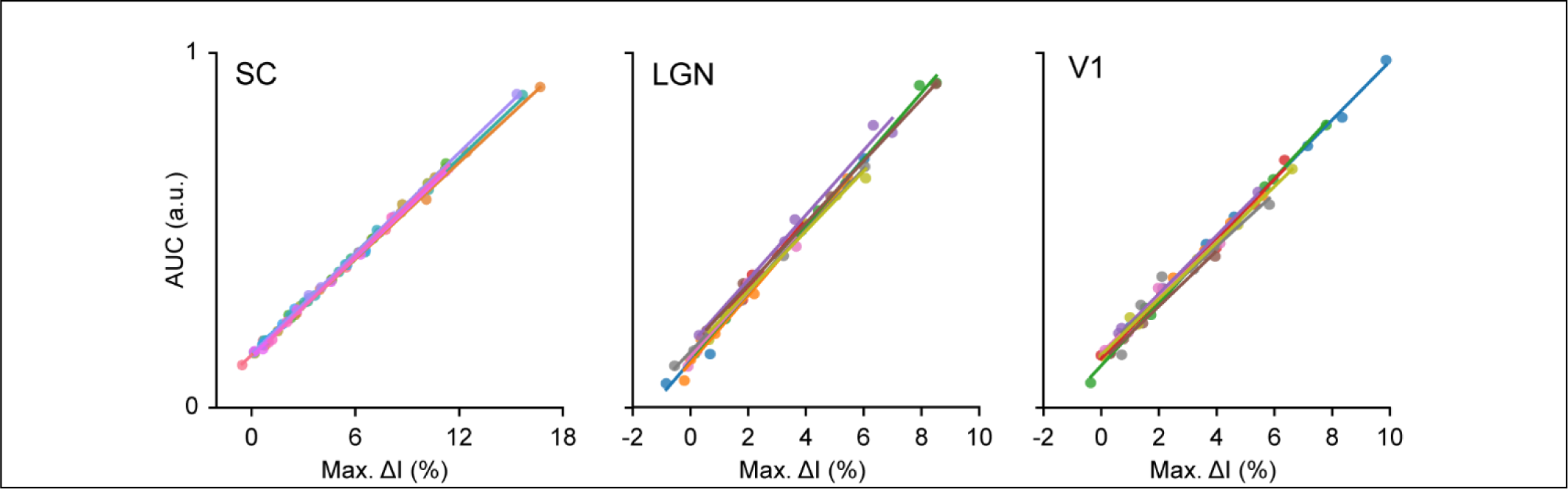
Correlation between maximum ΔI and area under the curve. Maximum of fUS signal variation (max. ΔI in %) with respect to the area under the curve of fUS signal variation (AUC in arbitrary units a.u.). Color corresponds to a recording session and dot represents a contrast condition.

**Figure S6.**
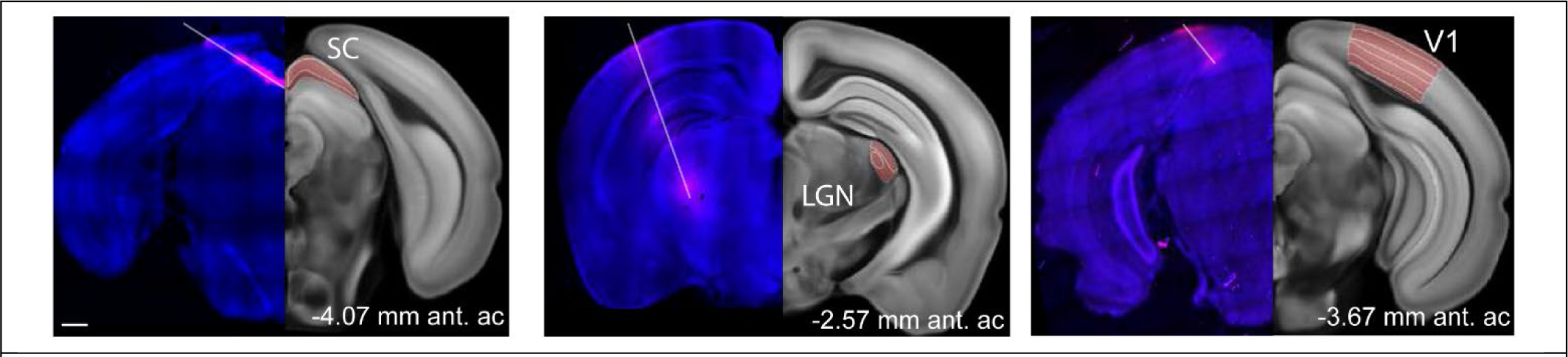
Neuropixels probe trajectories through the mouse brain. Histological reconstruction of the Neuropixels probe trajectories (white lines) showing DAPI staining (blue) and the fluorescent indicator DiI (pink) used to coat the probe. Trajectories are in the superior colliculus, lateral geniculate nucleus, or primary visual area; from left to right. Scale bar: 0.6mm.

**Table S1.** Mice and sessions.

| Mouse ID | Number of sessions per region |  |  |
| --- | --- | --- | --- |
|  | SC | LGN | V1 |
| 1 | 2 |  |  |
| 2 | 1 |  |  |
| 3 | 2 |  |  |
| 4 | 1 |  |  |
| 5 | 1 | 2 |  |
| 6 | 1 | 1 | 1 |
| 7 | 1 | 1 | 2 |
| 8 | 3 | 1 | 3 |
| 9 |  | 1 |  |
| 10 |  | 2 | 1 |
| 11 |  | 1 |  |
| 12 |  |  | 2 |
| Total | 12 | 9 | 9 |

**Table S2.** Coordinates used for Neuropixels probe insertion. CCS: Caudal confluence of Sinuses.

| Regions of interest | Position relative to CCS (mm) | Lateral position from midline (mm) | Angle (relative to horizontal) | Insertion depth (mm) |
| --- | --- | --- | --- | --- |
| SC | +0.5 | 2.5 | 25° | 3.0 |
| V1 | +0.4 | 3.0 | 12° | 3.5 |
| LGN | +1.6 | 3.6 | 60° | 3.5 |

